# Latent-centric Isotropic Resolution Enhancement for Expansion Microscopy Imaging via Neural Compression and Self-supervised Learning

**DOI:** 10.64898/2026.02.09.704755

**Authors:** Pin-Hsun Lian, Tzu-Yi Chuang, Ya-Ding Liu, Li-An Chu, Sheng-Cheng Chang, Yu-Chen Kuo, Wei-Kun Chang, Ann-Shyn Chiang, Gary Han Chang

## Abstract

Expansion microscopy (ExM) enables nanoscale imaging for disease characterization. However, whole-organ analyses remain limited by several challenges. Current super-resolution methods either require high-resolution ground-truth data or assume spatially uniform point spread functions—assumptions that rarely hold in whole-organ imaging with depth-varying aberrations and illumination drift. Existing methods also worsen storage demands by inflating already multi-terabyte datasets without using neural compression.

We propose a single-stage, self-supervised framework that addresses both resolution anisotropy and storage constraints through compression-aware isotropic super-resolution. Our approach combines a 2D lateral encoder that operates directly on raw slices to avoid memory limits with a lightweight volumetric decoder that preserves cross-slice continuity. A vector-quantized variational autoencoder (VQ-VAE) provides an information-sufficient bottleneck, achieving up to 128× slice compression and up to 8× axial resolution enhancement. This latent-centric design yields approximately 1000× reduction in storage compared with storing fully isotropic volumes.

The framework achieves higher GPU throughput, lower memory usage, and stronger multi-GPU scalability than prior methods. By designating compressed latent space as the native storage format, it enables efficient on-demand isotropic reconstruction directly from compact representations. This combination of isotropic enhancement and neural compression framework therefore makes large-scale, whole-organ ExM analysis practical while maintaining analysis-ready accessibility, addressing a bottleneck in translating ExM to clinical biomarker discovery.

## INTRODUCTION

Expansion microscopy (ExM, [1,2]) has emerged as a powerful tool for studying human disease pathophysiology, enabling nanoscale imaging across clinical specimens, revealing hidden cellular structures and disease markers [3,4], and extending to centimeter-scale imaging for whole-organ characterization that may inform future biomarker discovery [5]. However, scaling to whole-organ analyses is hindered by anisotropic resolution, caused by long-working-distance, detection optics for low numerical aperture, and light-sheet thickness trade-offs at large fields of view [6–8]. To achieve rapid, wide-field of view (FOV) acquisition, systems adopt low numerical aperture (NA) optics to increase working distance, which enlarges slice spacing and through-plane blur [9,10]. These constraints yield volumes with high lateral–axial anisotropy, distorting axial features and biasing quantitative 3D analysis [11,12]. Supervised deep learning-based super-resolution is widely used to mitigate axial anisotropy by learning mappings from anisotropic to near-isotropic volumes and restoring axial detail relative to high-resolution lateral references [13–15]. However, performance depends on available high-resolution lateral ground truth and often degrades under domain shifts across instruments and specimens. Moreover, naïve resolution enhancement inflates data volumes, whereas organ-scale ExM produces multi-terabyte datasets; therefore, methods that recover isotropy while maintaining compact, accessible representations are needed [16,17].

Self-supervised and unsupervised methods avoid the need for lateral ground truth and have gained interest for large-scale imaging [13]. However, many approaches rely on point spread function (PSF) modeling or uniform implicit blur assumptions, whereas depth-dependent PSFs, aberrations, motion, and illumination drift are common in whole-organ imaging [18,19]. Cyclic-consistent or two-stage frameworks enforce content consistency via dual generators with cycle-consistency loss [20] or a separately learned degradation model that approximates axial blurring. Physics-guided variants reduce to a single generator, but their stability depends on the accuracy and spatial invariance of the blur kernel. For example, a zero-shot deconvolution model is particularly sensitive to PSF mismatch and may introduce artifacts or provide limited gains when the assumed PSF is inaccurate [21]. Self-Net requires explicitly learning of the degradation function, after which a supervised DeblurNet is trained for isotropic recovery [22]. Alternative strategies, including saliency constraints [23] and fine-tuning with additional synthetic datasets of varied degradations [24], have been used to preserve overall structure during super-resolution.

Although self-supervised models can be effective, they cause increased computational cost, training instability, and reduced accuracy under depth-dependent degradations. System-level constraints further complicate implementation: full 3D generators are memory-intensive, whereas purely 2D slice-wise models compromise through-plane continuity and often require specialized post-processing to assemble coherent 3D volumes [25,26]. Even when super-resolution achieves near-isotropy, it often exacerbates ExM’s storage burden by ignoring neural compression advances. Evidence from compression models, including implicit neural fields, shows that large-scale biomedical datasets can be compressed with minimal reconstruction loss [27]. Neural compression for electron microscopy has also preserved ultrastructural details critical for connectomics while reducing storage by up to approximately 17-fold [28].

Recent advances in variational generative models provide a strong foundation. Variational autoencoders (VAEs) generate probabilistic latent representations well suited for compression [29]. Across EM datasets, VAEs deliver large storage reductions with minimal loss of task fidelity and report up to approximately 128× compression [30]. When trained on large domain-specific datasets, VAEs preserve features needed for clinical tasks [31]. VAEs can jointly optimize rate–distortion, outperforming traditional codecs and earlier neural methods on medical image compression [32]. Furthermore, vector-quantized VAEs (VQ-VAEs, [33]) discretize latent space via a learned codebook, mitigating posterior collapse, stabilizing training, and producing compact, indexable codes well suited to downstream tasks. VQ-VAEs have embedded clinically meaningful features, supporting compression and analysis of whole-slide pathology [34]. Recently, a VQ-VAE–transformer was scaled to population-level 3D brain MRI, generating morphology-preserving volumes that outperformed generative adversarial networks (GANs) on fidelity metrics, showing that codebook latents can remain compact while supporting large-scale quantitative analysis [35].

These challenges and opportunities motivate a different approach to isotropic enhancement for ExM—one that (1) restores 3D structure from anisotropic acquisitions without assuming a uniform PSF, (2) preserves axial continuity while avoiding the memory overhead of full 3D generators, and (3) integrates neural compression as a native component rather than a post-processing step. Such a system should produce compact, information-sufficient representations that serve both as the persistent storage format and as the substrate for on-demand 3D reconstruction.

Therefore, in this study, we propose a single-stage, self-supervised, compression-aware isotropic super-resolution framework that encodes anisotropic volumes into compact latent representations and reconstructs isotropic 3D volumes on demand. Unlike interpolation-first pipelines that rely on heavy 3D U-Nets or slice-wise 2D models, the proposed latent-centric design compresses first and reconstructs on demand, improving cross-slice continuity, scaling to organ-level data, and unifying compression with isotropic enhancement. A 2D lateral encoder operates directly on raw slices, avoiding volumetric upsampling and associated memory limits, whereas a lightweight volumetric decoder synthesizes a 3D volume preserving cross-slice continuity. A VQ-VAE generator provides an information-sufficient bottleneck. On representative sub-terabyte ExM datasets, the approach achieves up to 8× axial resolution enhancement and approximately 128× slice compression, yielding ≈10³× storage reduction relative to fully isotropic volumes. These properties deliver substantial storage savings and scalability, making on-demand isotropic enhancement feasible for whole-organ imaging in biomarker discovery.

## MATERIAL AND METHODS

### Model and data preparation overview

We propose a self-supervised variational encoder–decoder that operates directly on raw 2D lateral slices, compressing them into a compact, detail-preserving latent space sufficient for representing the 3D structure needed for isotropic, high-resolution reconstruction (**Fig. 1a**).

**Fig. 1.**
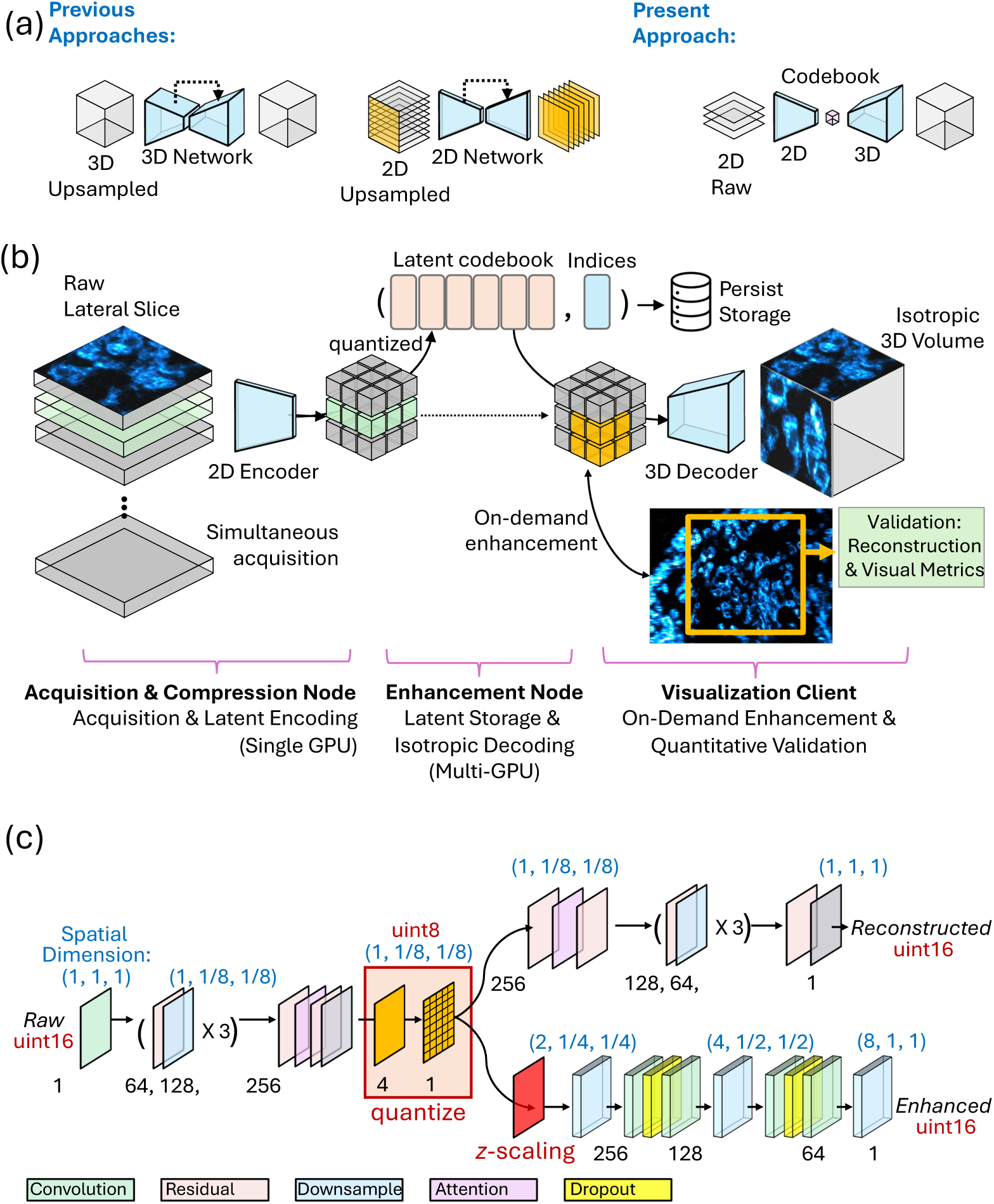
Latent-centric, on-demand isotropic 3D enhancement from 2D microscopy. (a) Raw 2D lateral slices are encoded into a compact vector-quantized latent and decoded directly to isotropic 3D image, in contrast to prior models that operate on upsampled 3D volumes or 2D slices. (b) System workflow: the acquisition node encodes and stores only codebook indices; an enhancement node decodes ROI on demand; a client visualizes and validates reconstructions. (c) Architecture: a 2D encoder produces quantized latents used by paired 2D and 3D decoders, with the 3D branch lifting along z-axis to achieve isotropy whereas the 2D branch validates the image reconstructed from the latent.

The model is trained with a combination of reconstruction and perceptual losses and, in the vector-quantized variant, a codebook and commitment loss that discretize the latent space into an informative code space. A lightweight 3D decoder expands the compressed latent space to synthesize an isotropic volume, aligning axial content with high-resolution lateral statistics via multi-view adversarial training and a patch-wise contrastive objective. Regarding the compute node, the compressed latent representation persists as a chunked Zarr [36] hierarchy aligned to the original acquisition, whereas the isotropic volumes are synthesized on demand for client-selected regions of interest and kept in memory for visualization, thereby avoiding persistent storage amplification and reducing the I/O overhead for downstream analysis (**Fig. 1b**).

We validated our approach across five biological datasets spanning multiple species, expansion factors, and imaging modalities: (A) *Drosophila* tyrosine hydroxylase (TH) neurons, 50× expansion microscopy (ExM), imaged with a large-FOV light-sheet microscope (LSM; NA = 0.137); (B) *Drosophila* TH neurons, 10× ExM, imaged with a spinning-disk confocal; (C) mouse brain blood vessels imaged with commercial LSM (SmartSPIM); (D) *Drosophila* neurons, 10× ExM, imaged with super-resolution microscopy of Airyscan; and (E) human surgical brain tissue, 10× ExM, imaged with a large-FOV LSM. Sample-preparation protocols were tailored to each specimen type. All human studies received IRB approval with informed consent. This multi-species, multi-modality validation demonstrates the broad applicability of our compression-aware isotropic enhancement framework across biological systems, expansion factors (10×–50×), and imaging platforms. Details of the biological samples, imaging and expansion protocols, as well as data dimensions used for model training and validation are provided in **Online Resource 1.**

### Self-supervised variational generative model for isotropic super-resolution and latent compression

The model accepts raw 2D lateral slices as input. In contrast to alternative approaches, these slices are not artificially upsampled, as the raw acquisitions already contain the requisite structural information. Upsampling at this stage would increase computational and memory demands without adding information. Instead, the framework directly encodes the slices into a compact latent representation, optimized with pixelwise reconstruction and perceptual losses. In the vector-quantized variant, an additional codebook loss ensures that discrete embeddings remain representative while preserving information.

Let *x* denote the input image and *x̂* denote the reconstructed image. The VAE is optimized with a pixelwise loss, ℓ*_p_* = |*x* − *x̂*|, for image reconstruction and a perceptual loss, ℓ_1_ = LPIPS(*x*, *x̂*) [37], for reconstruction fidelity. In the quantized model, a codebook loss ℓ*_cp_* is also included to enforce representation of discrete features.

Let *z_e_*(*x*) be the encoder output and *e_k_* the selected codebook entry. The quantization process can then be written as follows:

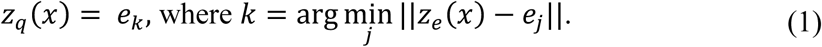

The codebook loss can be expressed as:

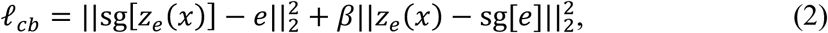

where the first term optimizes codebook embeddings *e* toward encoder outputs *z*_%_(*x*), the stop-gradient operator sg[·] prevents gradient backflow, and the second term encourages encoder commitment to selected codebook features, with ⋲ as the commitment weight. For a standard VAE, the model is regularized by the Kullback–Leibler divergence between the approximate posterior *q*(*z*|*x*) = *N*(μ(*x*), σ^2^(*x*)) and the Gaussian prior *p*(*z*) = *N*(0, *I*):

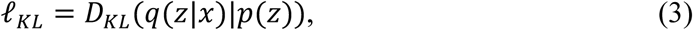

where μ(*x*) and σ^)^(*x*) are the encoder-predicted mean and variance, respectively. The encoder follows a hierarchical downsampling architecture similar to that used in the VQ-VAE and latent diffusion models [33]. After an initial convolution, residual blocks with self-attention progressively transform and downsample across multiple resolutions. The bottleneck combines two ResNet blocks for local feature refinement and an attention block for global context. The encoder output is mapped via a 1×1 convolution to the quantizer space and discretized by nearest-neighbor lookup in the codebook. The decoder mirrors this structure in reverse, progressively upsampling with residual blocks and optional attention until the original resolution is restored.

A lightweight 3D decoder expands the encoded latent space into fully isotropic representations with uniform voxel dimensions through self-supervised learning, eliminating the need for paired ground-truth data. The enhanced volume is optimized by using a multi-view adversarial loss, which helps align axial planes with high-resolution lateral imagery, and a patch-based contrastive (PatchNCE) loss [38], which preserves cross-plane content by maximizing agreement between corresponding structures while suppressing adversarial hallucinations and geometric drift.

In the multi-view adversarial formulation, two independent 2D PatchGAN discriminators (*D_yz_* and *D_xz_*) operating on the orthogonal *xz* and *yz* views are trained to distinguish the original high-resolution lateral slices from the enhanced outputs via an adversarial learning strategy as follows:

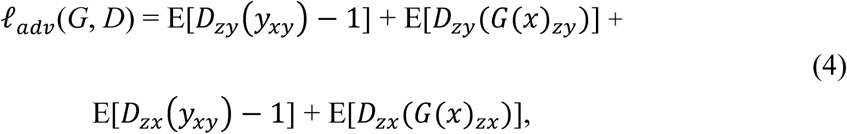

Complementarily, PatchNCE constrains the enhancement to remain faithful to the input by applying a patch-level InfoNCE objective between matched regions. For each anchor patch p_2_in the lateral view and its positive counterpart in the enhanced output, the objective increases the similarity of the positive pair relative to a batch of negative patches (q_2_), thereby maximizing mutual information across views and scales and mitigating the artifacts commonly associated with adversarial training. This process is expressed as:

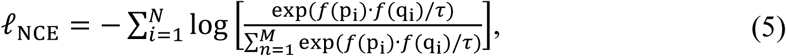

where *f*(·) is the encoder at the 2D patch level, *N* is the number of evaluated anchor-positive pairs, and τ is a temperature parameter.

Finally, a maximum intensity projection (MIP) loss is used to compare the original degraded images with the MIPs sampled at ω-voxel intervals along the axial dimension, preserving large-scale morphology while allowing the synthesis of high-frequency information between the axial slices.

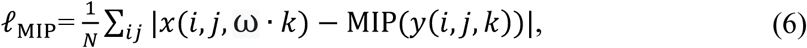

where MIP(·) denotes the maximum intensity projection.

The complete training objective is represented as:

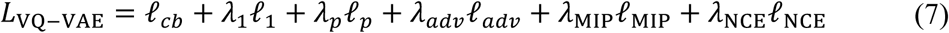

where ℓ*_cp_* is replaced with ℓ*_KL_* for the -quantized VAE. Further model architecture and training details are supplemented in **Online Resource 2**.

### Latent codebook and on-demand enhancement of isotropic microscopy volumes

The proposed compression-then-enhance system enables interactive exploration of large-scale microscopy datasets by synthesizing isotropic reconstructions on demand while maintaining compact latent representations. At the acquisition site, a dedicated compute node takes raw lateral slices as real-time input and executes the encoder on mini-batches sized to accelerator memory, processing one stack at a time as needed.

The encoder outputs latent representations written directly to persistent storage as chunked Zarr arrays aligned with the scan axes, with encoding scheduled to overlap with acquisition to optimize computational time and I/O. The system minimizes host–device transfers by streaming only the requested latent tiles to accelerator memory, where isotropic enhancement is performed end-to-end. This design reduces I/O stalls and avoids repeated host–device transfers, enabling practical latency for sub-terabyte to terabyte-scale volumes.

The latent representation persists as a Zarr hierarchy that mirrors raw data chunking, with vector-quantized variants storing per-location codebook indices and the latent features. Appropriate lossless or near-lossless compression is applied. Zarr metadata records voxel spacing, acquisition parameters, model identifiers, and mappings between regions of interest (ROIs) and latent tiles for reproducibility and downstream use.

Decoding, isotropic enhancement, and postprocessing are combined in a unified inference pipeline to avoid interstage communication overheads, and the models are executed in reduced precision (FP16) mode with calibration to preserve numerical stability. In the vector-quantized variants, the encoder outputs are discretized via a nearest-neighbor lookup strategy with a learned codebook, isotropic enhancement, and post-processing integrated in a unified inference pipeline to avoid interstage communication overhead. Models run in reduced precision (FP16) mode with calibration to preserve numerical stability. In vector-quantized variants, encoder outputs are discretized via nearest-neighbor lookup with a learned codebook, yielding compact 8-bit indices per spatial location.

During inference, ROI are mapped to the minimal covering set of latent tiles, with halo regions prefetched to reduce I/O latency and provide context for convolutional receptive fields. Reconstruction proceeds in a sliding-window manner over overlapping axial slabs (16 pixels), with tapered blending across overlaps to ensure continuity. Latent loading and enhancement overlap on separate computational streams, with synchronization limited to tile boundaries.

For large ROIs or high-concurrency scenarios, tiles are distributed across multiple accelerators and aggregated via peer-to-peer interconnections. Enhanced ROIs are retained in a memory-resident cache with time-to-live and least-recently-used eviction, enabling sub-second rendering. Chunks are streamed for visualization through Napari. The client interface supports real-time volume shading, orthogonal projections, and synchronized overlays of raw and enhanced data.

### Model training details

Training begins with sub-volumes of size 32×128×128 (z, x, y) uniformly sampled from the input microscopy volume. In-plane rotations are applied for augmentation. The architecture uses a fully convolutional design and supports inference on larger fields of view. The VQ-VAE encoder–decoder uses a base channel width of 128 and channel multipliers of 1, 2, 2, and 4, with one residual block per scale level. The latent representation has four channels, vector-quantized with a codebook of 256 entries of four dimensions each. Input volumes are encoded into latent space, quantized by nearest-neighbor lookup in the codebook, and reconstructed by the decoder. Training uses pixel-level reconstruction and perceptual similarity losses, along with codebook and commitment penalties. The quantization term is weighted by 1.0, and reconstruction combines L1 distance with LPIPS. This phase stabilizes the latent space and ensures sufficient representational diversity.

The 3D decoder is trained simultaneously in a self-supervised manner to achieve axial super-resolution. The generator is optimized with a weighted sum of adversarial, pixel reconstruction, and contrastive losses. Contrastive regularization uses a PatchNCE formulation: from each crop, 256 patch features are sampled at intermediate generator layers. For each location, a positive pair is formed from aligned input and reconstruction features, whereas negatives are drawn from other positions in the same sub-volume. This enforces local content preservation without large batch sizes. Optimization uses an initial learning rate of 2×10⁻⁴, decayed with a cosine annealing schedule over 2,000 epochs. Training is conducted with batch size 1 to minimize the memory footprint when using consumer grade GPU.

### Result Validation and quality measurements

Validation uses perceptual and structural metrics computed within randomly sampled ROIs. Primary perceptual distances include FID, KID, and LPIPS, where lower values indicate closer agreement. Baselines are defined as discrepancies between raw lateral and raw axial views; principal comparisons are between raw lateral and enhanced axial reconstructions. Comparisons include two representative models. OT-CycleGAN, an extension of CycleGAN with optimal transport regularization to preserve mass and geometry between axial and lateral planes, serving as a strong unpaired generative baseline. ZS-DeconvNet, a zero-shot deconvolution approach that estimates dataset-specific blur on the fly and performs non-blind deblurring without paired supervision, serving as a classical deconvolution comparator.

I/O throughput, per-kernel execution time, and end-to-end ROI latency are recorded under representative loading during system-level validation to ensure interactive performance. Downstream utility is assessed by running segmentation models on reconstructed ROIs and reporting IoU and F1 relative to manual annotation. Model and codebook identifiers are embedded in file metadata to enable deterministic rollback or retraining when drift detectors signal changes in the optical system, illumination profile, or specimen domain. By persisting only discrete or low-bit-rate latent representations and synthesizing isotropy on demand via an accelerator-resident, stream-overlapping pipeline, the framework maintains plane-aware perceptual fidelity and practical wall-clock latency for large-scale microscopy while avoiding prohibitive storage expansion and I/O bottlenecks.

## RESULTS

### Isotropic enhancement reduces axial–lateral disparity across species and modalities

To assess whether the self-supervised model restores axial fidelity and reduces axial–lateral disparity across diverse biological structures and imaging modalities, we first evaluated human brain surgical tissue imaged with a customized mesoscope at 50× ExM (voxel size 1.88 × 1.88 × 15.04 µm) and Drosophila brain TH (tyrosine hydroxylase, TH)-positive neurons imaged with confocal microscopy at 10× ExM (voxel size 1 × 1 × 8 µm). The model was applied to raw anisotropic stacks, and the enhanced axial views were compared against the raw lateral views as reference. In both datasets with distinct structures and optical properties, the enhanced axial views recovered fine neuronal details, and disparities between raw axial and lateral views were substantially reduced after isotropic enhancement (**Fig. 2a, b)**. Previously conflated features in the raw axial views became separable, with axial line intensity profiles exhibiting sharper peaks; full width at half maximum (FWHM) decreased substantially (0.93 to 0.27 and 41.2 to 12.8 µm for human and Drosophila tissue, respectively. Fourier analysis indicated reduced anisotropy, with axial-to-lateral high-frequency power ratios improving from 7.4 to 1.2 and from 3.05 to 0.99 for human and Drosophila tissue, respectively (**Fig. 2c, d**).

**Fig. 2.**
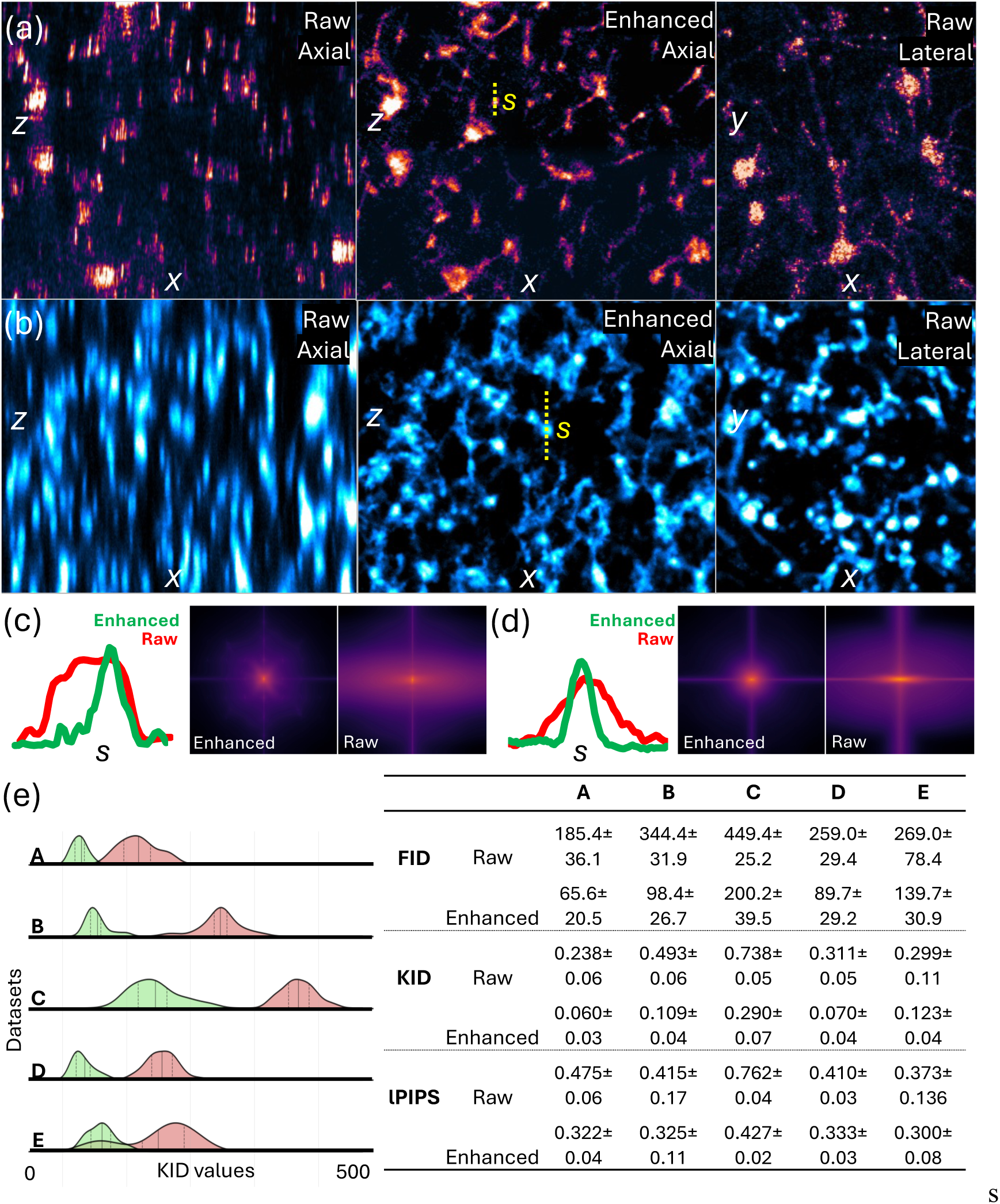
Isotropic resolution enhancement increases axial resolution and reduces anisotropy. Raw axial, enhanced axial, and native lateral views for human brain surgical tissue (a) and Drosophila TH neurons (b). Regions where features appearing conflated in the raw axial view are separable in the enhanced axial view. FWHM and spectrum analysis show significant signal improvement on the enhanced axial view of the (c) human brain and (d) Drosophila TH⁺ neurons, respectively. (e) Cross-dataset summary (A–E) of FID, KID, and LPIPS between axial views and raw lateral references. Dataset mapping: A = Drosophila TH neurons, 50× expansion microscopy (ExM), large FOV light-sheet microscopy (LSM); B = Drosophila TH, 10× ExM, spinning disk confocal; C = mouse vasculature, commercial LSM (SmartSPIM); D = Drosophila neurons, 10× ExM, super-resolution microscopy (Airyscan); E = human brain tissue, 10× ExM, large FOV LSM.

We further quantified enhancement performance across species and modalities by computing FID, KID, and LPIPS between the enhanced axial and native lateral views. Compared with baseline values (raw axial vs. lateral), enhanced images showed substantial improvements across five datasets, indicating consistent gains across diverse structures and systems (**Fig. 2e**). Together, these results show that the model improves axial fidelity and enables isotropic 3D reconstructions from confocal to high-throughput mesoscope acquisitions, where throughput is prioritized over axial sectioning.

### Comparative validation against orthogonal references and self-supervised baselines and robustness against real-world degradations

To validate structural fidelity, we compared enhanced axial slices from large-FOV LSM volumes with an anisotropic ratio originally of 1:8 (lateral:axial resolution of 1.88:15.04 µm) with lateral slices acquired from the orthogonal direction as the reference.The enhanced images successfully recovered tubular morphologies and suppressed axial streaking that fused structures in the baseline (**Fig. 3a**). Furthermore, we acquired high-NA mouse brain images with lower anisotropy (original lateral:axial resolution of 1.8:4.0 µm) and downsampled along the axial axis to create a volume also with an anisotropic ratio of 1:8 (lateral:axial resolution of 0.5:4.0 µm), using the original high-resolution views as reference. The enhanced axial views sharply delineated capillary boundaries and accurately restored centerlines, closely matching the lateral reference (**Fig. 3b**). These results indicate that the model generates 3D reconstructions that agree with orthogonal reference views as well as the reference from the original high-NA stacks.

**Fig. 3.**
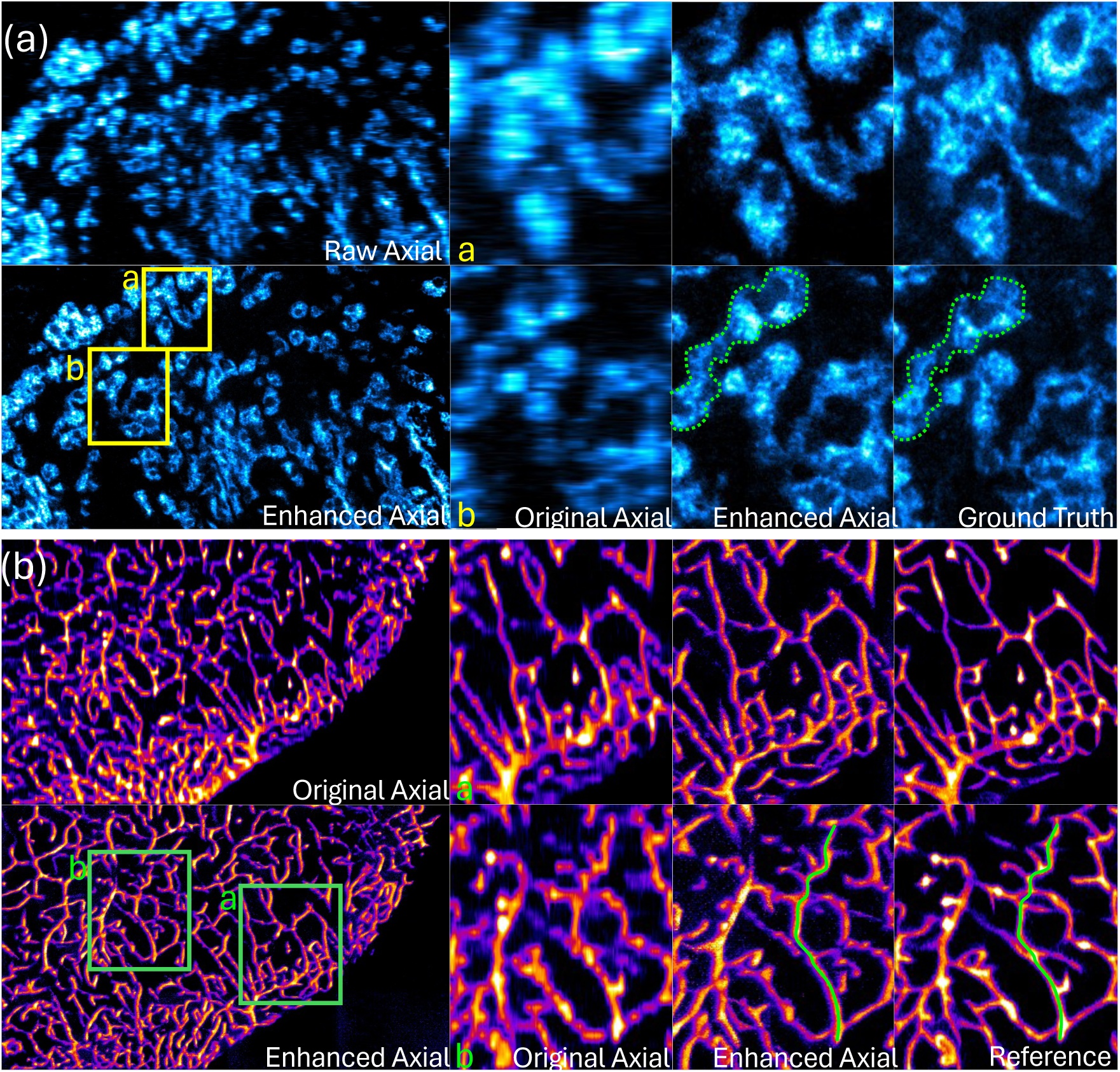
Ground truth validation of structural fidelity by reference of orthogonal acquisition and high NA image. (a) Drosophila neuron image volume with anisotropy of 1:8: raw axial, enhanced axial, and reference orthogonal acquisition. (b) Downsampled mouse brain vasculature image volume with anisotropy of 1:8 with the original high-resolution image as the reference: downsampled axial, enhanced axial, and original high-resolution view.

To evaluate robustness under real-world degradations not captured by PSF models, we compared our method against other self-supervised approaches. Applied to Drosophila neuron volumes acquired with Airyscan—prone to slice-to-slice lateral drift due to short-working-distance optics—the model reconstructs isotropic 3D structures without drift-specific training (**Fig. 4a**). During side-by-side comparisons with ZS-DeconvNet and OT-CycleGAN, our method preserved lateral-like morphology with fewer discontinuities under both lateral drift and high axial anisotropy from rapid acquisition (**Fig. 4b**). We further tested robustness under synthetic depth-dependent degradations (lateral drift, intermittent slice loss, depth-dependent SNR gradient, and light scattering) that were absent during training. The model generalized to these conditions, producing axial views with perceptual quality and morphology closely matching the native lateral scans (**Fig. 4c**). Additional details of ground-truth image validation and robustness against real-world degradation conditions are further provided in **Online Resource 3.**

**Fig. 4.**
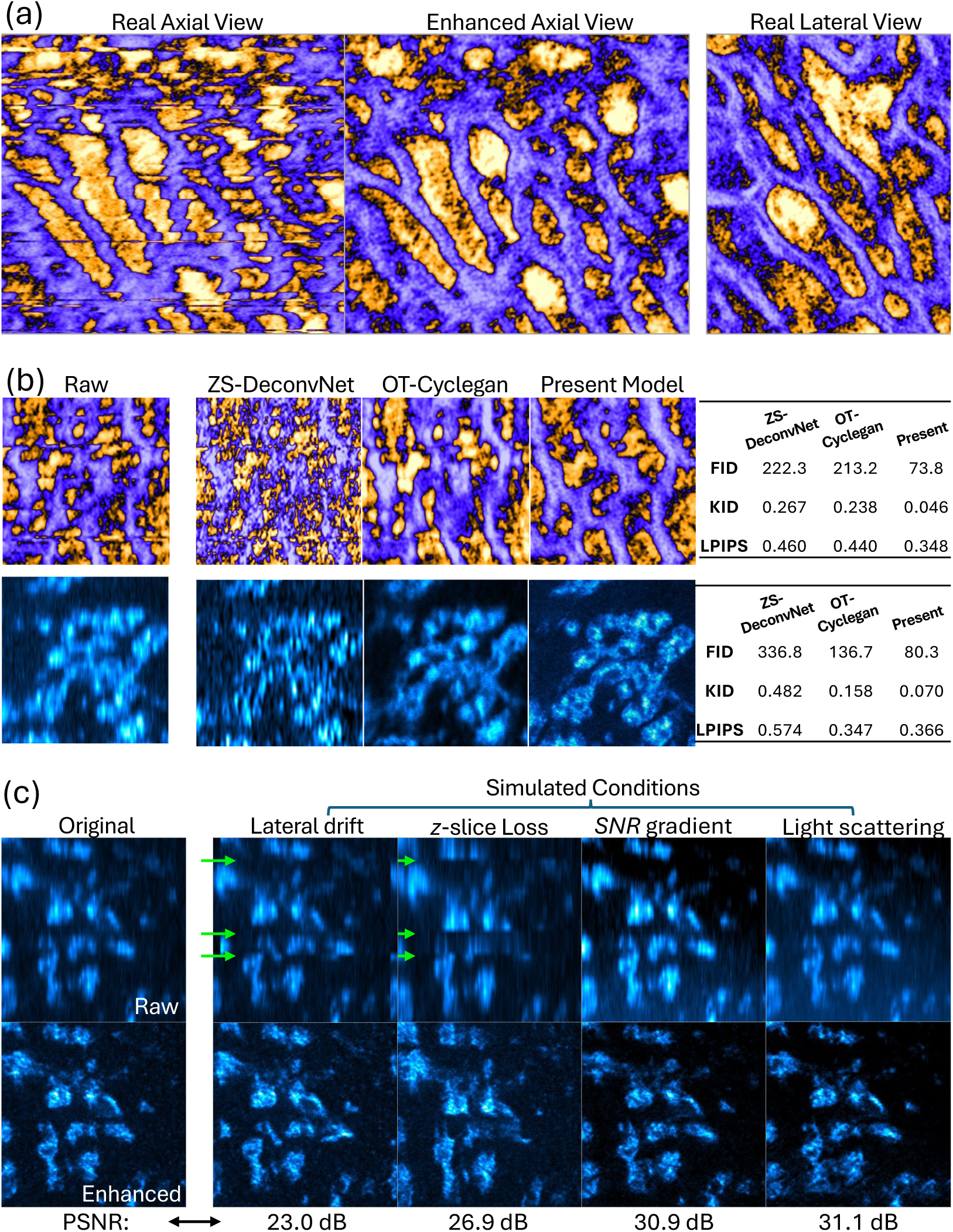
Robustness to depth-dependent degradations across acquisition conditions. (a) The model generates high-quality axial view from Airyscan imaging affected by slice-to-slice lateral drift. (b) Comparative study of different self-supervised models in isotropic enhancement under lateral drift (top row) and high axial anisotropy from mesoscope rapid acquisition (bottom row): ZS-DeconvNet, OT-CycleGAN, and the proposed model. (c) Generalization to unseen depth-dependent degradations: lateral drift, random z-slice loss, depth-dependent SNR gradient, and light scattering.

### Unsupervised 3D segmentation by stochastic sampling of the generative model

We investigated whether the enhanced axial view could support zero-shot cell segmentation. We generated standard deviation (Std) maps from multiple stochastic forward passes. From the enhanced axial view *x*, candidate binary masks were defined via thresholding *m* = 1{ x > τ }, and the Std map was computed across *K* stochastic reconstructions (**Fig. 5a**). This procedure enhanced diffuse fluorescence boundaries into uncertainty-bounded contours, transforming soft edges into sharp delineations An ablation on stochastic samples showed boundary convergence at *K* ≈ 20, indicating that only modest sampling is required for robust contour extraction (**Fig. 5b**). We tested zero-shot instance segmentation by combining isotropic enhancement with cell segmentation foundation model of CellSAM [39] using the large-FOV LSM Drosphilla data. Although raw anisotropic images exhibited elongated cell edges along the axial axis, enhanced images with uncertainty maps showed refined boundaries and improved instance separation when comparing to manual ground truth (**Fig. 5c**). Furthermore, the enhanced axial view, while assisted with uncertainty map, delineated neuronal morphologies from uneven fluorescence signals in the human brain tissue dataset, matching manual annotations (**Fig. 5d**).

**Fig. 5.**
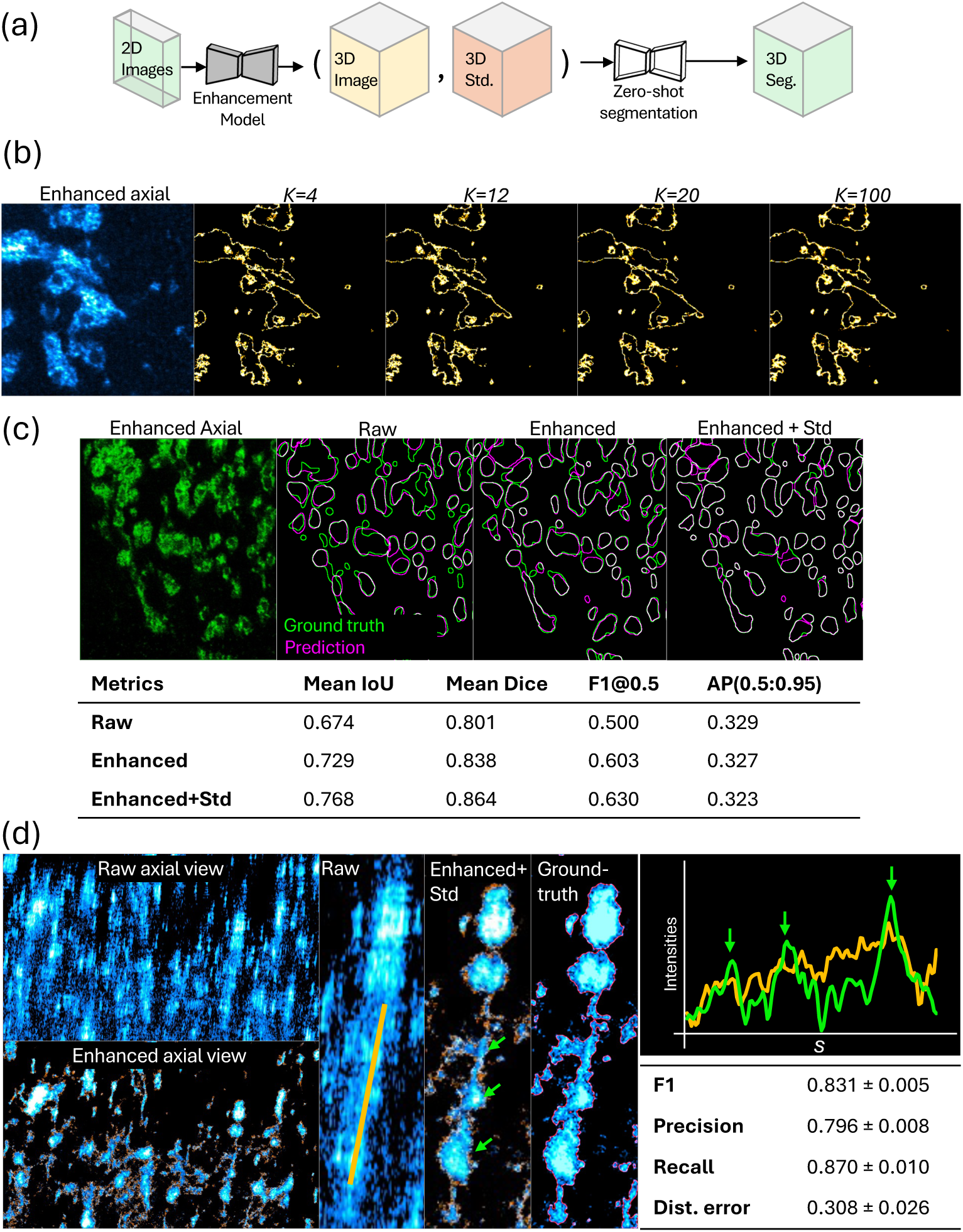
Zero-shot cell segmentation using enhanced axial views with uncertainty quantification. (a) Standard deviation maps generated from K stochastic forward passes creates sharp, uncertainty-bounded contours, faciliating zero-shot segmentation. (b) Boundary convergence analysis shows optimal performance at K≈20 iterations. (c) CellSAM foundation model applied to enhanced Drosophila LSM data demonstrates improved instance separation compared to raw anisotropic inputs when validated by instance segmentation metrics against manual ground truth. (d) Enhanced axial views with uncertainty maps successfully delineate neuronal morphologies in human brain tissue, matching manual annotations as evaluated by boundary segmentation metrics.

### Consistent multi-sample neuron–protein colocalization across heterogeneous acquisitions

To assess generalizability to multi-sample, multi-structure analyses, we tested whether isotropic enhancement enables reliable neuron–protein colocalization across brains with heterogeneous acquisitions. Using available 2D lateral segmentations, we concatenated these annotations with raw lateral images as multi-channel inputs; the model then produced isotropic 3D reconstructions and corresponding segmentations in a single pass (**Fig. 6a**). Applied to simultaneously acquired dorsal paired medial (DPM) neurons and vesicular monoamine transporter (VMAT) volumes [40], enhancement corrected axial elongation that had merged neuronal and protein structures. Co-localization events that appeared as single overlaps in raw axial views resolved into multiple isotropic appositions after enhancement, reducing false positives and improving quantification (**Fig. 6b**).

**Fig. 6.**
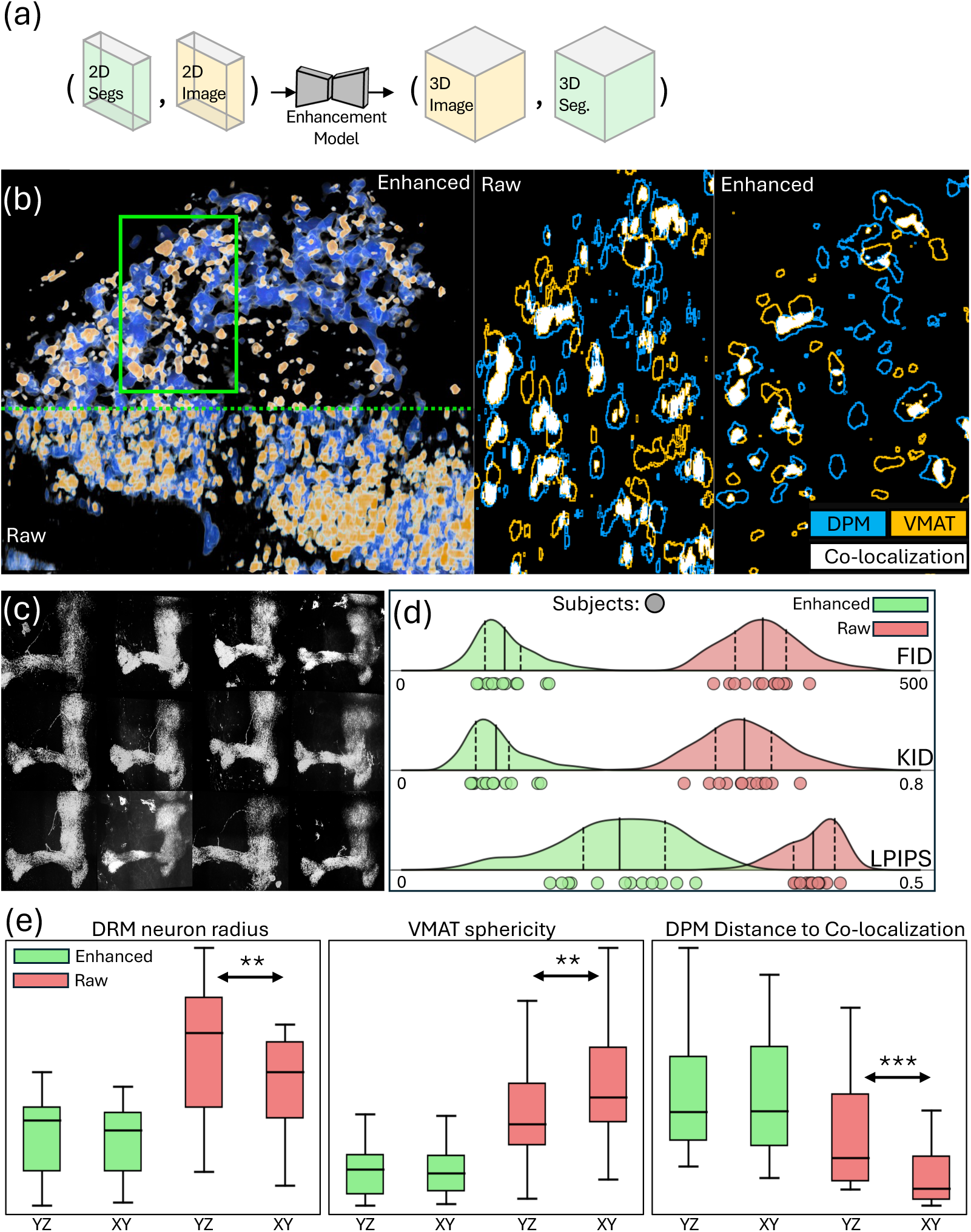
Multi-sample neuron–protein colocalization enabled by isotropic enhancement and single-pass 3D segmentation. (a) Raw lateral images and their 2D segmentations are concatenated as multi-channel inputs, the model outputs isotropic 3D reconstructions and corresponding 3D segmentations in a single pass. (b) Example volumes of DPM neurons and VMAT: raw axial, enhanced axial, and native lateral views, showing separation of overlaps observed in the raw axial view. (c) Visualization across heterogeneous fluorescence conditions (n = 12 samples). (d) Distribution of image-quality metrics between axial and lateral views across n = 12 samples. (e) Morphology and colocalization metrics across samples: DPM radius, VMAT sphericity, and DPM–VMAT colocalization distance for raw axial, native lateral, and enhanced axial views. Statistical significance indicated by asterisks: * p < 0.05, ** p < 0.01 (*t*-test). DPM: dorsal paired medial neuron and VMAT: vesicular monoamine transporter.

The method was robust across heterogeneous fluorescence conditions. Across all samples (n = 12; **Fig. 6c**), enhanced axial views achieved isotropic resolution, confirmed by distribution-level image quality metrics (FID, KID and LPIPS; **Fig. 6d)**. Morphology metrics—including radius of DPM neurons, sphericity of VMAT, and DPM–VMAT colocalization distance—showed that discrepancies between raw axial and lateral measurements were eliminated after enhancement. Measurements from enhanced axial views matched those from native high-resolution lateral view across all samples (**Fig. 6e)**. Thus, the proposed model could consistently reconstruct biologically faithful 3D morphologies and improve the quantification of interaction information across multi-sample datasets without retraining under heterogenous conditions.

### Sub-terabytes scale computational study

We next evaluated computational scaling. For representative 102 GB anisotropic volumes divided into non-overlapping 384³-voxel patches, encoding achieved real-time compression in 44.5 min on a single NVIDIA A6000 GPU—faster than the microscopy acquisition time, enabling deployment at acquisition nodes for immediate compression during imaging sessions. Decoding scaled near-linearly with GPU count: total decode time decreased from 10,481 s on a single GPU to 2,612 s on four GPUs, whereas per-patch time remained constant at approximately 1.91 s. Overall reconstruction time was instead bottlenecked by file serialization, which remained constant at approximately 5,400 s regardless of GPU count. This performance profile supports a two-stage deployment: single-GPU encoding nodes for real-time compression during acquisition, followed by batch super-resolution reconstruction on multi-GPU systems for downstream analysis.

**Table 1.** Encoding (compression) and decoding (super-resolution) performance reported as per-cube calculation time and total runtime.

| Process | Number of GPUs | GPU computing time | CPU computing time | Serialization time | Total runtime |
| --- | --- | --- | --- | --- | --- |
| Encode | 1 | 0.209 s | 0.297 s | 0.008 s | 2,672 s |
| Decode | 1 | 1.915 s | 0.104 s | 0.931 s | 13,458 s |
| Decode | 2 | 1.910 s | 0.098 s | 0.942 s | 8,419 s |
| Decode | 4 | 1.909 s | 0.105 s | 0.996 s | 6,040 s |

## DISCUSSION

Whole-organ expansion microscopy faces major challenges from anisotropic resolution and massive storage demands that block translation to clinical biomarker discovery. We developed a unified framework that addresses both issues through compression-aware isotropic reconstruction. We show that a single-stage, self-supervised VQ-VAE achieves up to 8× axial resolution enhancement while compressing data approximately 128×, yielding an effective approximately 1000× storage reduction. The method generalizes across microscopy modalities and biological specimens, with the largest benefits for high-throughput systems where speed compromises axial detail. These findings establish that variational latent representations can serve as both the native storage format and the substrate for on-demand isotropic reconstruction, making large-scale ExM analysis practically feasible for clinical applications.

Without multi-stage training or multi-generator architectures, our VAE-based model enhances highly anisotropic inputs to isotropic quality, matching the performance of leading self-supervised methods. Unlike U-Net–style models that use extensive skip connections to pass multi-scale details directly to the output, our approach enforces a compact, information-sufficient latent space via a variational bottleneck. This enables downstream tasks to run directly on compressed latent codes rather than on full-resolution volumes. To our knowledge, few prior isotropic enhancement methods explicitly exploit a shared compressed latent representation. Given rapid advances in neural compression for microscopy, this workflow represents a scalable solution. Because the latent representation faithfully reconstructs raw images, it serves as the native substrate for downstream tasks.

Our method simplifies training and reduces computational complexity compared to approaches that learn separate degradation models. This is a key advantage, since many self-supervised baselines require retraining for different structures or imaging modalities. Our approach also remains robust to spatially varying degradations, as shown by experiments under diverse simulated conditions and a real-world case of sampling drift in super-resolution microscopy. By avoiding the need for explicit PSF characterization or synthetic training data, our method sidesteps the impracticality of exhaustively modeling all degradation conditions. Instead, it leverages the consistent appearance statistics of high-resolution views. This provides a complementary path to physics-informed super-resolution methods that explicitly model forward degradations. Furthermore, uncertainty estimates in our approach identify regions where self-supervision is sufficient and where additional measurement-consistency regularization may be required.

We acknowledge several limitations. Despite validation across diverse modalities and against orthogonal views from the same system used as pseudo–ground truth, exact voxel-wise correspondence remains difficult to quantify. ExM samples are not rigid, and deformations during sequential acquisitions can misalign structures; such misalignments are magnified at organ scale. Thus, our evaluation emphasizes regional features and perceptual image quality metrics rather than voxel-to-voxel comparisons. Although mature dual-view systems are inefficient for large-scale organ imaging, they can provide better-aligned ground-truth data for future validation. In addition, although our method does not require PSF calibration and demonstrates robustness to moderate spatially varying degradations, extreme aberrations or rapidly varying PSFs may still degrade performance and should be monitored during training and validation. Finally, although single-stage training and latent-centric inference reduce computational burden, we still observe significant non-GPU overhead from I/O, tiling, and stitching, which pose challenges for very large datasets. Although we implemented multi-GPU data-parallel inference, future work could explore shared-memory inference and rendering strategies to enable seamless whole-organ visualization and reconstruction.

## CONCLUSION

We present a single-stage, self-supervised framework that addresses the dual challenges of anisotropic resolution and massive storage that currently limit the potential of expansion microscopy in human disease research. By integrating VQ-VAE–based compression with isotropic super-resolution, our approach achieves up to 8× axial resolution enhancement and approximately 128× compression, producing an effective approximately 1000× reduction in storage requirements while generalizing across diverse imaging modalities and biological specimens. This latent-centric design removes critical technical barriers to centimeter-scale ExM imaging. By making large-scale ExM analysis computationally and economically feasible, this work accelerates the translation of ExM’s unique capabilities—revealing hidden cellular structures and disease markers in whole-organ characterization—toward practical clinical biomarker discovery.

## Notes

### Competing Interest Statement

The authors have declared no competing interest.

